# Spontaneous network coupling enables efficient task performance without local task-induced activations

**DOI:** 10.1101/2020.05.04.075804

**Authors:** Leslie Allaman, Anaïs Mottaz, Andreas Kleinschmidt, Adrian G. Guggisberg

**Affiliations:** Division of Neurorehabilitation, Department of Clinical Neurosciences, University Hospital of Geneva, Genève, GE, 1211, Switzerland; Division of Neurology, Department of Clinical Neurosciences, University Hospital of Geneva, Genève, GE, 1205, Switzerland

## Abstract

Neurobehavioral studies in humans have long concentrated on changes in local activity levels during repetitive executions of a task. Spontaneous neural coupling within extended networks has latterly been found to also influence performance. Here, we intend to uncover the underlying mechanisms and the interaction with task-induced activations. We demonstrate that high performers in visual perception and motor sequence tasks present an absence of classical task-induced activations, but, instead, strong spontaneous network coupling. Activations were thus a compensation mechanism needed only in subjects with lower spontaneous network interactions. This challenges classical models of neural processing and calls for new strategies in attempts to train and enhance performance.

## Introduction

The understanding of how the brain resolves basic behavioral problems such as sensory perception or motor planning has been one of the central questions of science. Over the last decades, neuroscience has accumulated evidence that the successful performance of behavioral tasks is associated with patterns of neural activations at specialized brain areas during the task. Activations can be observed with brain imaging as changes in a neural signal during the task as compared to a baseline signal which is typically measured in the absence of a specific task or stimulation, Figure 1A. When investigating field potentials induced by large neural assemblies, task-induced activations manifest as an increase of fast oscillations in the so-called high-γ frequency range (>60 Hz) and a reduction of slower rhythms (Crone *et al*., 2006). Since the first recordings of neural activity in humans, it is known that neural assemblies spontaneously produce prominent oscillations of an electromagnetic potential, in particular at a frequency of about 8 to 12 cycles per second even when the recorded subjects are at rest. These spontaneous oscillations, named *α*, are suppressed when subjects engage in a task (Pfurtscheller *et al*., 1996). Importantly, α suppression has been shown to improve stimulus perception (Ergenoglu *et al*., 2004; Babiloni *et al*., 2006; Hanslmayr *et al*., 2007), speed of detection (Thut *et al*., 2006), discrimination (van Dijk *et al*., 2008), as well as lateralized visuo- spatial attention (Thut *et al*., 2006; Romei *et al*., 2010). Therefore, the posterior α rhythm has for long been viewed as reflecting processes of attentional disengagement and being intrinsically inhibitory (Pfurtscheller *et al*., 1996; Klimesch *et al*., 2007; Jensen and Mazaheri, 2010). The suppression of α and β amplitudes, often referred to as *event-related desynchronization* (ERD), is a widely used index of brain activation (Pfurtscheller and Lopes da Silva, 1999).

**Figure 1.**
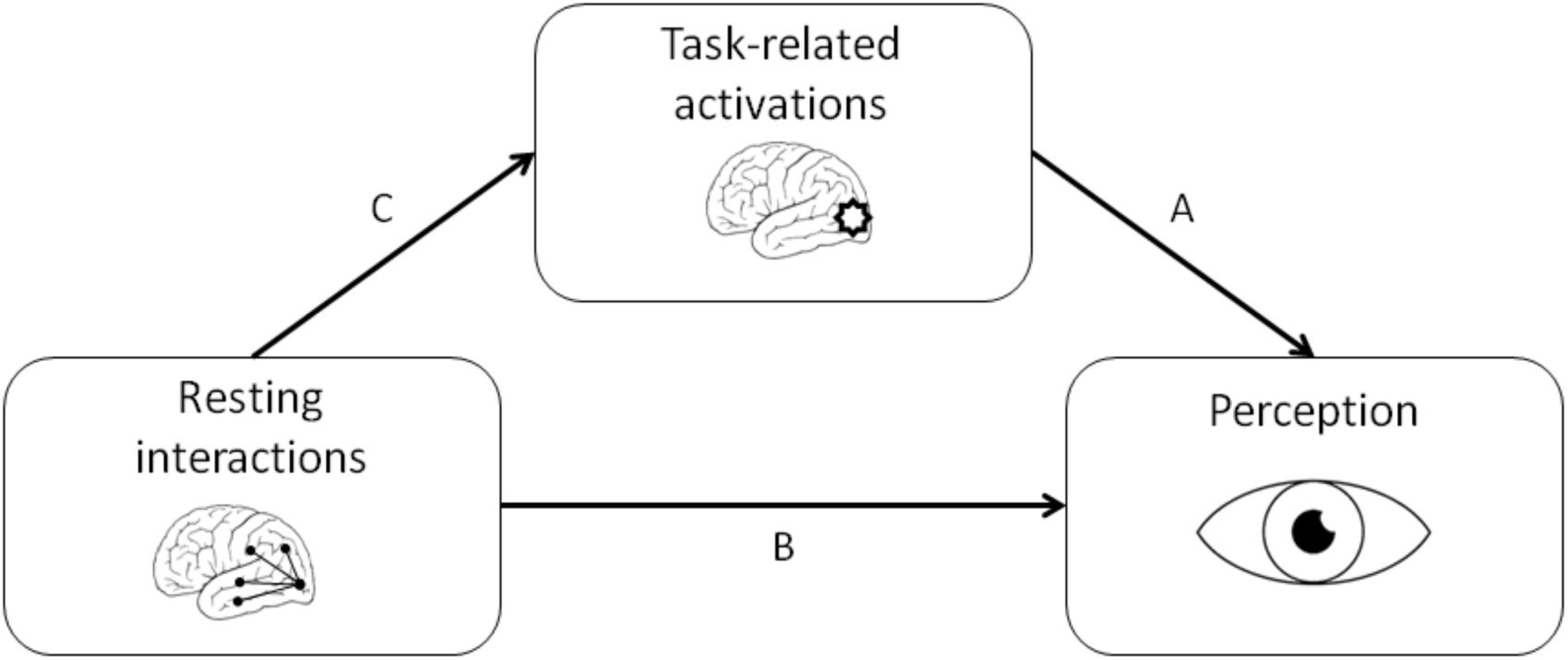
Schematic illustration of the relations between neural activity patterns and visual perception.

Nevertheless, more recent research showed that there is more to task-free baseline states and α oscillations than inhibition. A large body of research has demonstrated that spontaneous brain activity is organized into coherent networks (Greicius *et al*., 2009). Neural coupling (i.e., functional connectivity, FC) between brain areas states preceding a task facilitates perception and behavioral performance (Weisz *et al*., 2014; Sadaghiani *et al*., 2015; Strauß *et al*., 2015; Iemi *et al*., 2019), thus suggesting that network coupling plays an active role (Figure 1B). This also applies to longer, spontaneous, task-independent neural coupling, in particular in α frequencies, which is positively correlated with performance (Hipp *et al*., 2011*b*; Guggisberg *et al*., 2015). For instance, the more spontaneous α activity in Broca’s area is coherent with the rest of the brain, the better subjects are able to produce words (Guggisberg *et al*., 2015). Phase coupling of the α rhythm in visual areas can also predict the detection of a transient visual target (Hanslmayr *et al*., 2005). Patients with brain pathologies such as stroke and Alzheimer’s disease have reduced α-band coupling, and the degree of this reduction correlates linearly with the severity of the behavioral impairment in a variety of functions, such as language, motor function or spatial attention (Dubovik *et al*., 2012, 2013).

In sum, not only neural activations during tasks, but also spontaneous network interactions are associated with successful task performance. However, their respective role and the interactions between them (Figure 1 C) are unclear. In the present study, we thus compared the role of resting-state neural coupling (FC) and task-induced neural activations on behavior, within paradigms of visual perception and motor planning. We hypothesized that both neural patterns influence behavior. Our findings are in line with these hypotheses but go even further. Neural α-band coupling was found to be a more efficient neural mechanism underlying perception and motor planning than task-related activations, which were a compensation mechanism necessary only in subjects with baseline states of low neural coupling. This challenges the traditional notion of a primary importance of task-induced activations.

## Results

### Experiment 1

In a first experiment, twenty healthy subjects performed a perithreshold visual detection task interspersed with resting periods (see Figure 2). This enabled us to study the two central processing patterns of interest, classical local activations and spontaneous network interaction states, within a single paradigm. Subjects were presented with brief visual targets to the left, right, or bilateral visual field. For each subject, the contrasts were chosen such that about half of presentations were consciously perceived in each condition while the others were missed (at the same external stimulation parameters).

**Figure 2.**
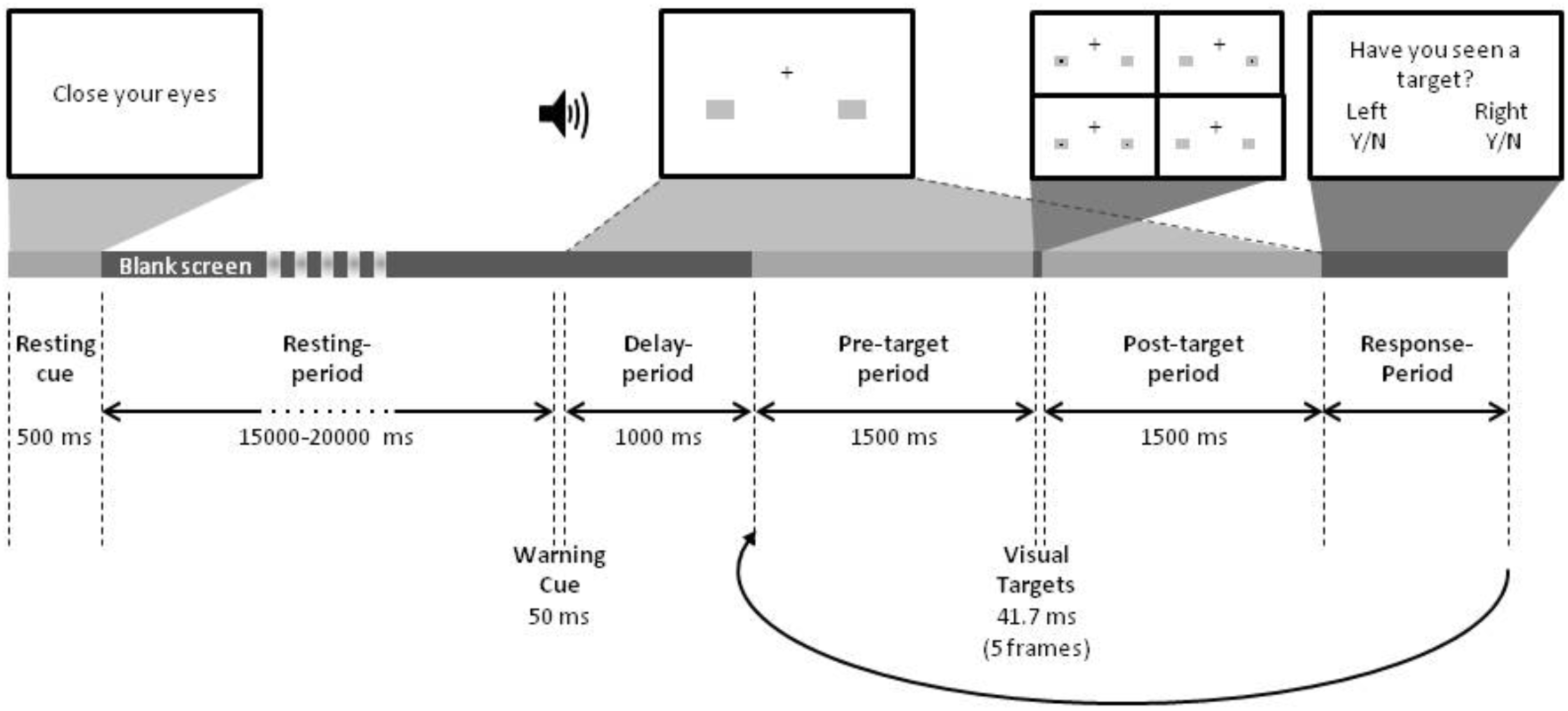
Experimental paradigm and time course of a single block. The task consisted of blocks starting with a resting period followed by seven trials of a target detection task (see Methods for details). For visualization purpose the background is shown here in white, whereas the background of the task was dark grey.

Figure 3 illustrates the behavioral results. False alarm rates were very low in all participants (range 0-4.2%, median=.4%) and Grier’s B’’ values ranged from .711 to 1 (median=.97), showing they all adopted a conservative response criteria as instructed. One sample t-tests showed that performances in left, right, and bilateral stimulation conditions did not differ from 50% accuracy (all p>.11), confirming that we were able to display perithreshold contrasts to our participants.

**Figure 3.**
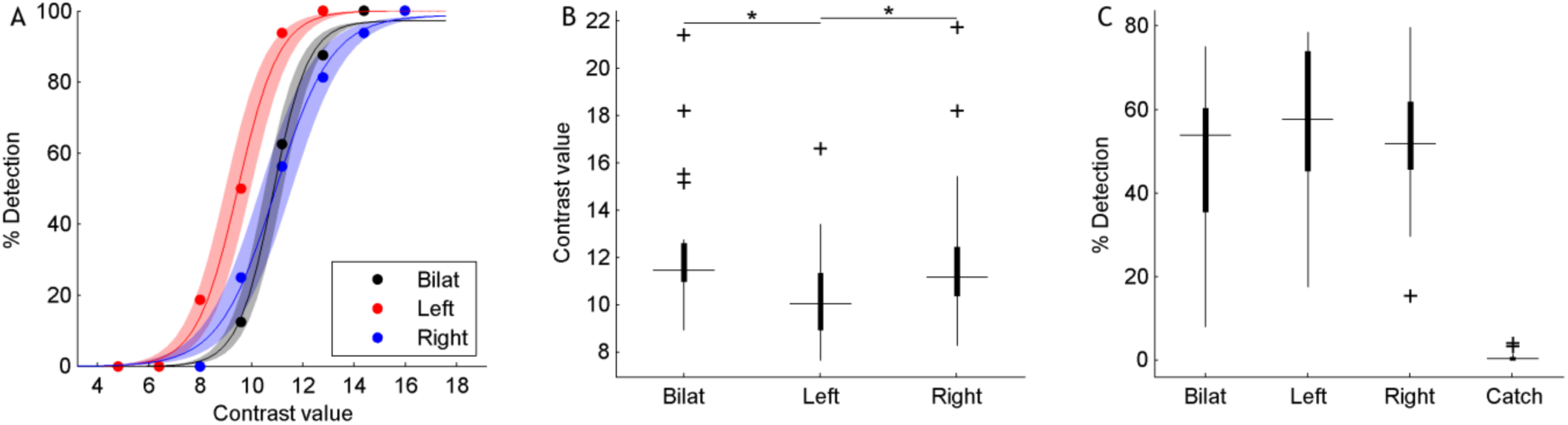
Behavioral results. Sensitivity curves for one subject (A), threshold contrast values (B) and detection rates (C) for all subjects and conditions. * = p < 0.05

We then used inverse solutions to reconstruct oscillations in the brain grey matter from high-density surface EEG and computed local oscillation power during the target detection task. In addition, we estimated spontaneous neural interactions during the inter-block resting states using the graph theoretical measure weighted node degree (WND), indicating the sum of FC between a brain area of interest and all other areas (Newman, 2004).

#### Task-induced activation and spontaneous network coupling

We first confirm that visual target presentation induced, on average, the classical α-band power decrease at bilateral visual areas after left as well as right target presentation as compared to a baseline just before target presentation (Fig. 4 A-B). When assessing all frequency-bands at an anatomically defined, bilateral, occipital region of interest (ROI, see Figure S2), one can observe the expected classical task-induced activation pattern, consisting of enhanced high-γ activity and suppressed activity at slower bands (Fig. 4 C). Since low frequency power decreases for left and right targets were bilateral (see Discussion), we henceforth report results based on pooled data from left and right conditions at the bilateral occipital ROI. Figs. S3 and S4 further demonstrate that the upcoming findings were consistent after left and right target presentation.

**Figure 4.**
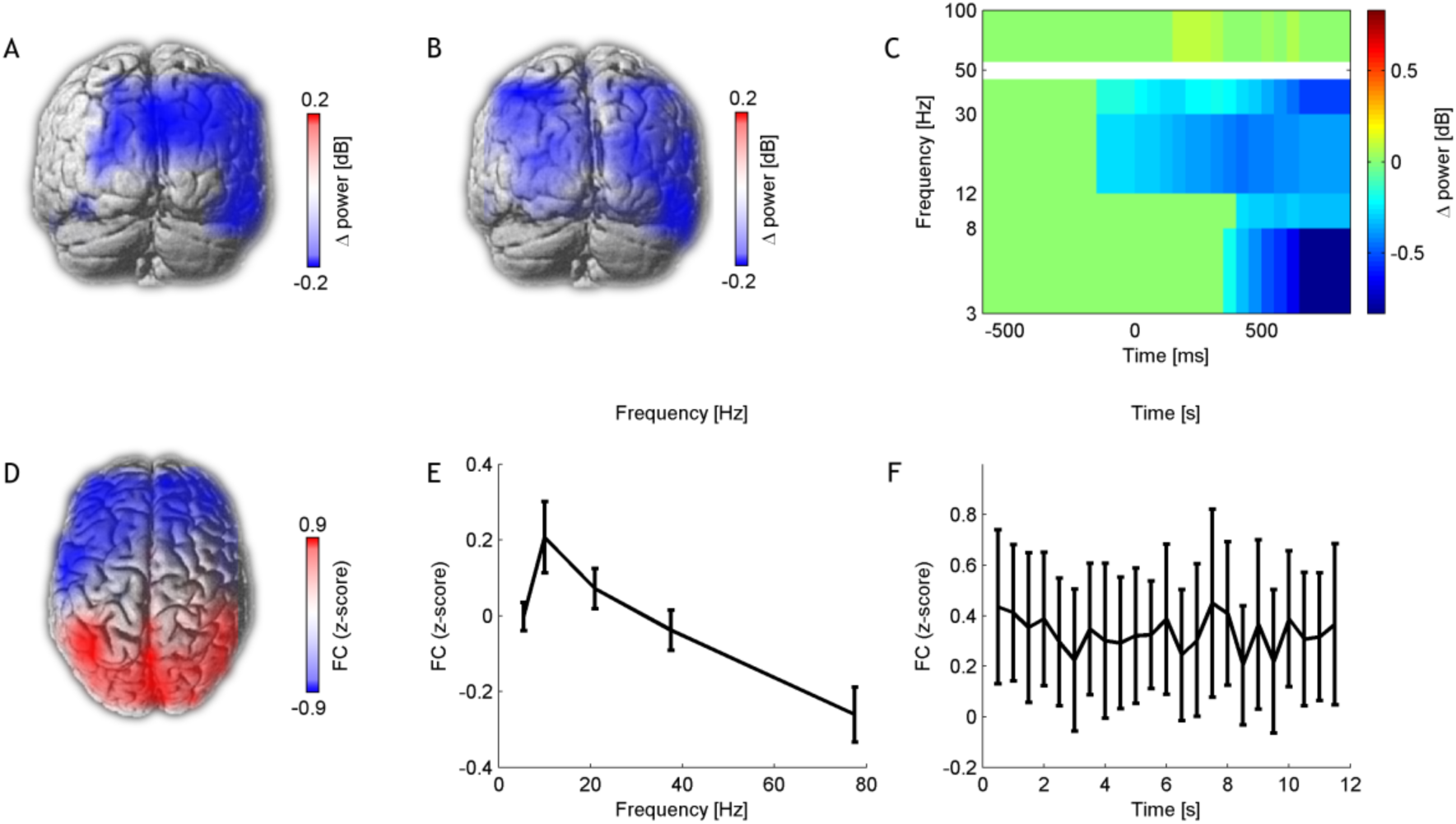
Characteristics of investigated neural processes. Average event-related in-task power changes induced by targets presented to the left (**A**) and right (**B**). Blue colors represent regions with significant power decrease in the α-band after the presentation of a target (p<0.05, 5% FDR corrected) on a 3D rendering of a template brain. **C** Time-frequency decomposition of power changes in a bilateral occipital ROI as compared to a baseline at 500 to 300 ms before target onset, as induced by a left or right visual stimulus (p<0.05, 5% FDR corrected). The topography of resting-state α-band functional connectivity shows an occipito-frontal gradient (**D**). Frequency spectrum of resting-state functional connectivity in a bilateral occipital ROI (**E**). **F** A-band functional connectivity variation over the inter-block resting-periods. Error bars indicate mean ± 95% confidence interval.

When assessing global resting-state FC, i.e., the WND, of each brain area, we reproduced the occipito-frontal gradient (Fig. 4 D) and predominance at α frequencies (Fig. 4 E) described previously (Guggisberg *et al*., 2008; Hillebrand *et al*., 2012). A repeated-measures one-way ANOVA of WND at the ROI revealed a significant effect of frequency band (F_4,76_=33.34, p<.001) with FC values significantly higher in the α band than in all other bands (p<0.05, Tukey-Kramer HSD). This confirms that the α band was the preferential frequency for neural communication (Chapeton *et al*., 2019). A repeated-measures one-way ANOVA showed no effect of time across the twelve seconds of inter-block resting- periods (F_22,418_=.56, p>.9), hence demonstrating that the α-band FC states were stable during resting periods (Fig. 4 F).

#### Event-related desynchronization and neural coupling both predict perception

As predicted in our first hypothesis, a within-subjects analysis of single trials confirmed that both, in-task power decrease and spontaneous α-band FC at the ROI are associated with better visual perception. Seen trials induced greater event-related power decrease in α, θ, and β-bands in the post-stimulus period than missed trials (p<0.05, FDR corrected), Fig. 5 A. Spontaneous FC was greater during resting periods before seen than before missed trials in the α frequency band (t_19_=3.03, p=.035, FDR corrected), but not in the other frequency bands (all uncorrected p>.05), Fig. 5 B. A multivariate, mixed-model ANOVA confirmed that both, α-band FC (F_1,7006_=6.8, p=0.009) and α-band power decrease from baseline (F_1,7006_=12.9, p=0.0003) had independent impact on perception at the single trial level. Importantly, unlike FC, local resting-state α power at the ROI did not predict perception (F_1,7008_=0.0, p=0.95).

**Figure 5.**
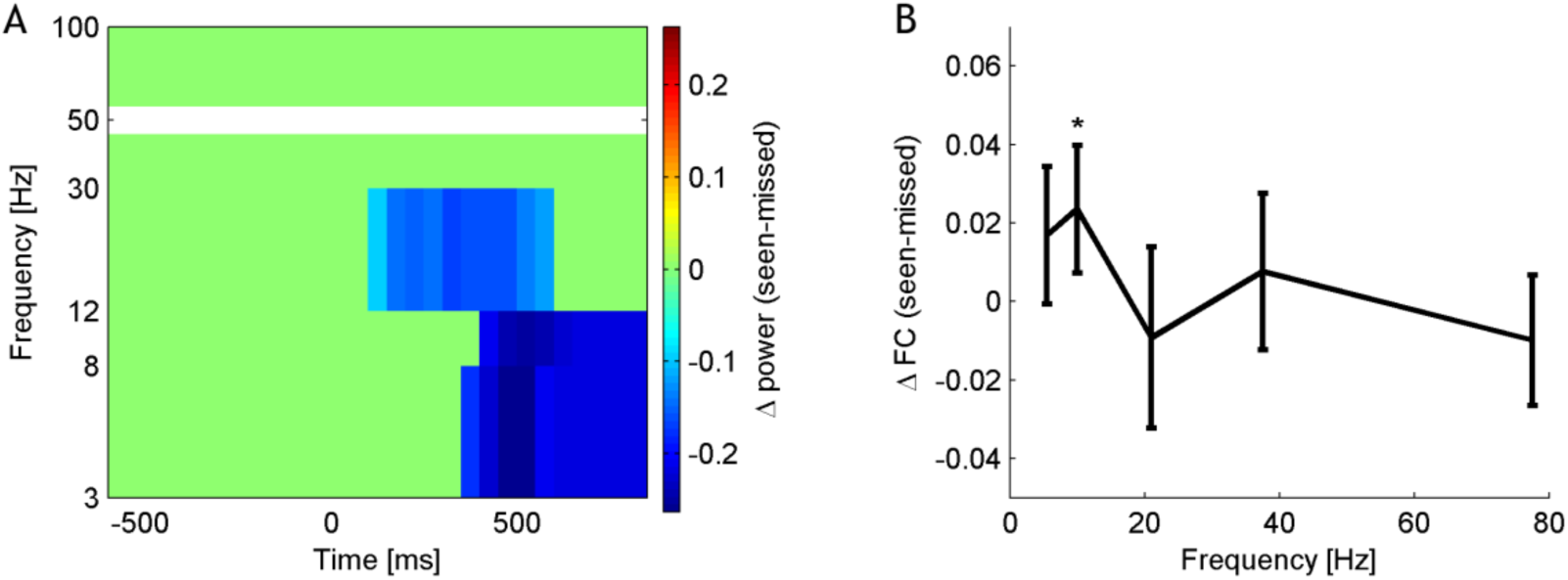
Neural processes predicting visual perception. Seen stimuli were associated, on average across all subjects, with greater power changes in θ, α, and β bands during the presentation of seen stimuli than missed trials (**A**, p<0.05, 5% FDR corrected), in accordance with the classical pattern of task-induced neural activations. We also observed greater α-band coherence between the region of interest and the entire cortex during resting periods before perceived stimuli than before missed trials (**B**, p=.035, 5% FDR corrected), in accordance with previous findings of a positive impact of α-band coherence during rest for task performance (Guggisberg *et al*., 2015). Error bars indicate mean ± 95% confidence interval.

#### ERD and FC have opposite effects on visual performance

We then subjected the neural patterns associated with visual perception identified in the previous screening steps to between-subject covariations between behavioral and neural levels. This confirmed that subjects with greater resting-state FC in the α-band had proportionally *better* visual perception percentage (r=.45, p= .045, Fig. 6 A). Importantly, this was not the case for local resting-state α power (r=.10, p=.66). Conversely, subjects with greater in-task power decrease in seen than missed trials in θ (r=.45, p=.045), α (r=0.37, p=0.10), and β bands (r=.47, p=.038) tended to perform proportionally *worse* in visual perception percentage (Fig. 6 B-D). Thus, classical local in-task activation was associated with worse task performance while abundant spontaneous interactions were present in subjects requiring particularly low contrasts for correct visual perception.

**Figure 6.**
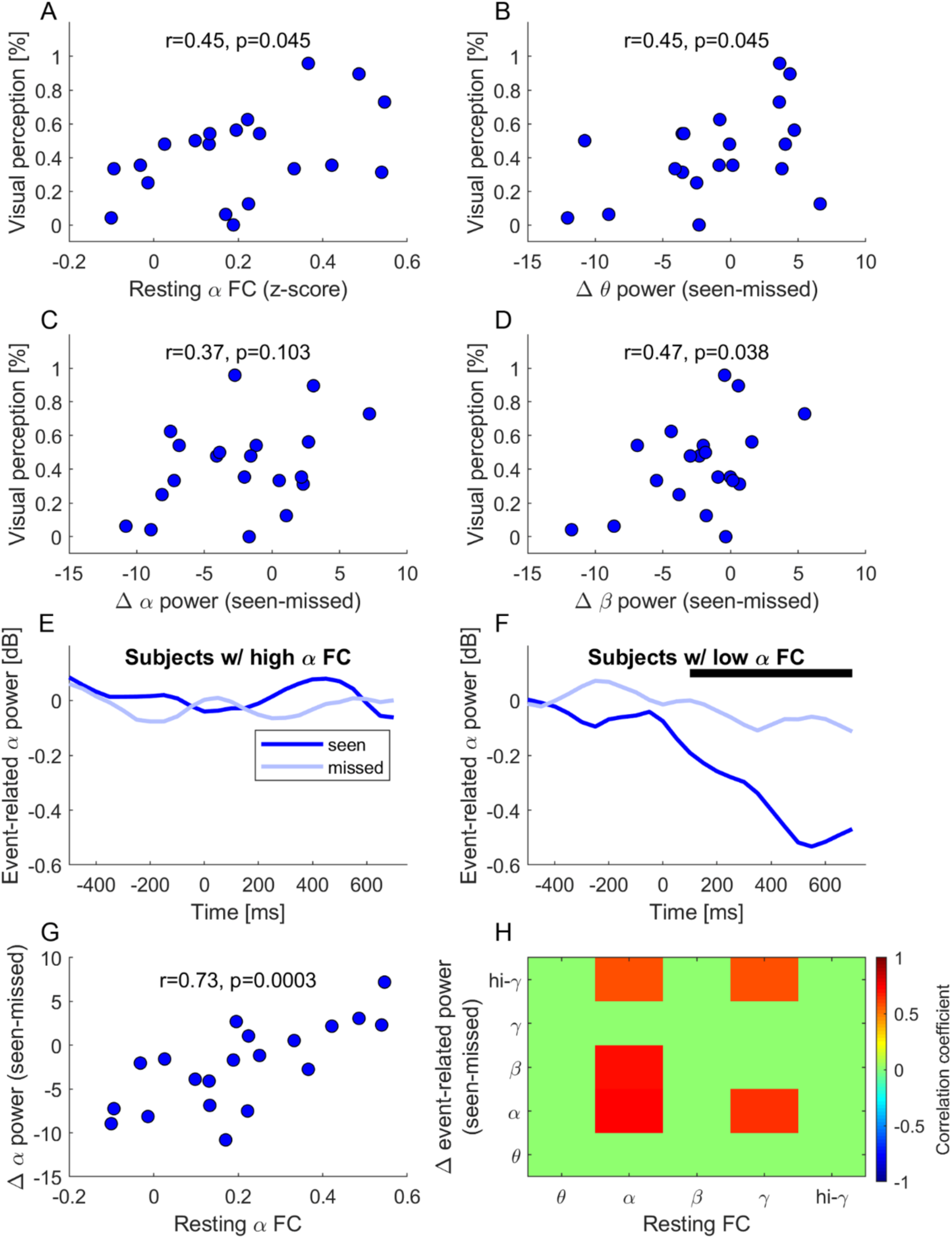
Between-subject co-variation of neural processing and target perception. Subjects with greater spontaneous α-band WND in the occipital ROI showed proportionally better visual detection rates (**A**). Conversely, subjects with greater in-task θ, α, and β power difference (seen-missed) tended to have proportionally lower detection rate (**B** - **D**). Dividing the sample in two equal groups based on their levels of resting-state α-FC (median cutoff) showed that only subjects with low α-band FC level demonstrate the classical task-induced power decrease (**E**). The black line indicates time windows in which the difference between seen and missed trials is significant (p < 0.05). Conversely, subjects with high α-band FC show an absence of α-power decrease following stimulus presentation, regardless of their awareness of the target (**F**). Association between resting-state α-band FC and in-task α power difference (**G**) and same association for all bands (Pearson correlations, p<0.05, 5% FDR corrected) (**H**).

#### Spontaneous network coupling reduces the need for in-task activation

To examine underlying reasons for the inverse impact of resting-state α-FC and in-task power differences on visual performance, we divided the participants in two equal groups based on their levels of resting-state α-FC (median cutoff). This showed that only subjects with low α-band FC demonstrate α-power decrease after target presentation (Fig. 6 E). In these subjects, a two-way repeated-measures ANOVA shows a main effect of time (F_25,459_=7.95, p<.001) and perception (F_1,459_=12.32, p=.007), suggesting significant power decrease over time which was greater during seen than during missed trials. Conversely, subjects with high spontaneous α-band FC actually showed an absence of α-power decrease following stimulus presentation, regardless of their awareness of the target (Fig. 6 F). A two-way ANOVA shows no effect of time (F_25,459_=.33, p>.99) or perception (F_1,459_=.74, p=.41), and no interaction (F_25,459_=1.07, p=.38). The same conclusion had to be drawn for an explorative analysis of in-task power in all other frequency bands, as no frequency band showed significant differences between seen and missed trials in participants with high spontaneous α-band FC (Fig. S5). Moreover, a voxel-wise analysis of the whole brain did not find any area with significant α power decrease from baseline (p>0.5, 5% FDR corrected) in this subject group.

Bi-variate correlation analyses between the two neural patterns further demonstrated that subjects with higher levels of α-FC needed proportionally less power decrease in seen trials compared to missed trials in α (r_18_=.73, p=.008, corrected, Fig. 6 G), as well as β (r_18_=.69, p=.009, corrected), and high γ bands (r_18_=.57, p=.047, corrected). An association was also found between γ resting-state FC and power difference in α (r_18_=.63, p=.026, corrected), and high-γ (r_18_=.57, p=.047, corrected). Crucially, unlike α FC, local α power during the resting periods (p>0.69, corrected), or FC in the other frequency bands did not correlate with in-task power differences in any band (Fig. 6 H, p>0.5, corrected). Thus, the correlation was not merely due to a trivial dependency between FC and power or to mathematical coupling of a difference value.

#### Network correlates of leftward spatial attention bias

A leftward visuo-spatial attention bias is commonly found in young healthy subjects (Thut *et al*., 2006; Learmonth *et al*., 2015). This was also the case in our paradigm. A Friedman test showed a significant effect of the condition on threshold contrast values (*χ*^2^_(2)_=27.70, p<.001), as threshold values for left targets were lower than for right and bilateral targets (p<0.05, Tukey-Kramer HSD). We assessed whether this behavioral bias was also reflected in spontaneous network interaction states, both on the single trial and the subject level. Top down control is thought to be mediated by right parietal areas (Gillebert *et al*., 2011). Accordingly, we defined the right intraparietal sulcus (Fig. 7 A) as ROI for these analyses (Eickhoff *et al*., 2005), using areas hIP1, hIP2 (Choi *et al*., 2006) and hIP3 (Scheperjans *et al*., 2008).

**Figure 7.**
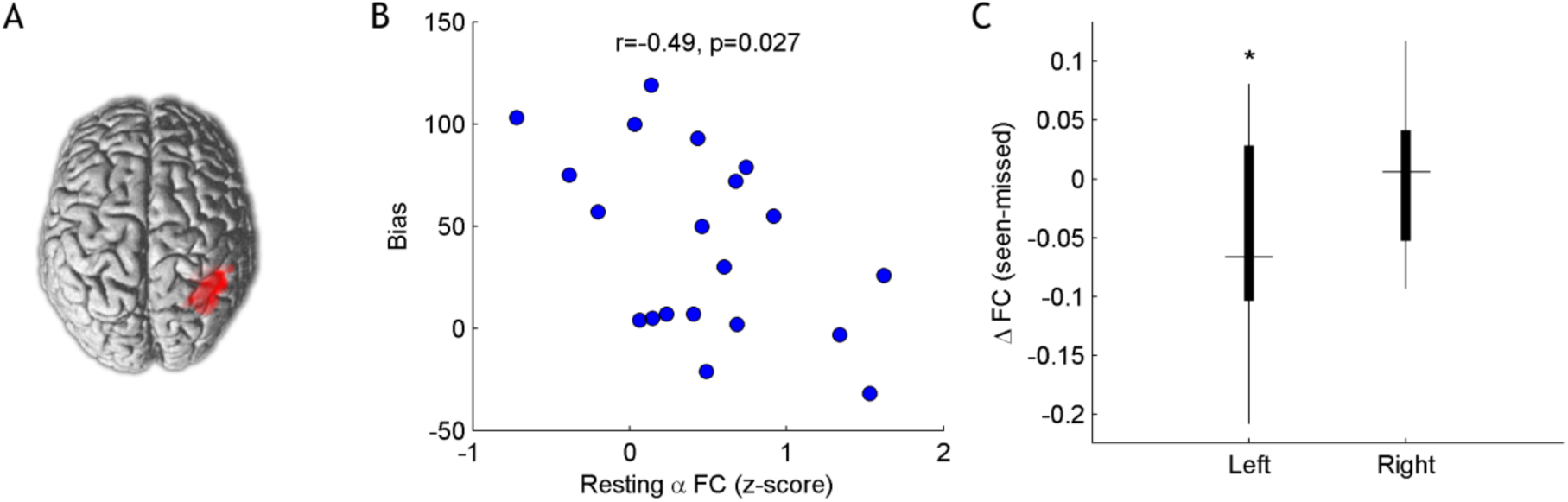
Spontaneous network correlates of leftward spatial attention bias. **A** Parietal region of interest (ROI). **B** Pearson correlation between resting-state α-WND of the parietal ROI and visuo-spatial leftward bias in bilateral stimulus presentations. **C** Difference in α-WND preceding unilateral stimulus presentations. * = p < 0.05, 5% FDR corrected.

As predicted, a negative correlation was found between resting-state α-FC in the parietal ROI and the visuo-spatial leftward bias (r=-.49, p=.027), see Figure 7 B. Difference in α-power during the task, in contrast, did not significantly correlate with the visuo-spatial leftward bias (r=-0.33, p=0.15). Regarding unilateral targets, one-sample t-tests showed that the difference in FC in the ROI before seen versus missed trials was significant for left targets but not for right targets (respectively p=.025 and p=.73, 5% FDR corrected), see Figure 7 C. The same analysis regarding power difference in the peri-stimulus period showed no significant difference (left: p=0.16, right: p=0.21, 5% FDR corrected).

These findings demonstrate that participants with greater α-band coupling in the right intraparietal sulcus overall showed less spatial bias towards the left visual field. On a trial-by-trial basis, this effect is apparent on unilateral targets, for which we show that greater coupling before stimulus presentation disfavors perception of left, but not right targets. We suggest that greater coupling of the parietal ROI allows a stronger correction of the spontaneous leftward attentional bias when the task requires bilaterally distributed attention. At the trial level, variations of said coupling mediate the overall advantage shown for left targets.

#### Inter-individual differences in FC

Given the prominent impact of spontaneous network coupling, we were interested in factors that might promote it. An explorative analysis did not reveal significant effects of age, gender, hours of sleep the night preceding the recording, or subjective fatigue during the task (p>0.29, 5% FDR corrected). The amount of pre-existing experience with visual discrimination might contribute to spontaneous neural coupling, but was not quantified in subjects of experiment 1.

### Experiment 2

To further clarify this point, we performed a second experiment. Twenty healthy participants performed a finger tapping task (see Methods) which is frequently used to assess motor skills (Zhang *et al*., 2012). Nine participants had previous experience with piano playing (median 12 years, range 1 to 12 years) and thus with motor sequence tasks, which enabled us to test the influence of previous experience with the task on FC. Experiment 2 further intended to demonstrate that our findings obtained with visual perception generalize to other task types.

The finger tapping task induced, on average, robust ERDs in θ, α, β, and γ bands during the motor planning phase in centro-parietal brain areas contralateral to the moved hand (Fig. 8 A) and in primary motor areas in particular (Fig. 8 B). ERDs in motor ROIs did not correlate with performance in this task in any frequency band (|r|<0.39, p>0.085). Conversely, when investigating spontaneous α-band coupling as reconstructed from 5 minutes of resting-state recording, we reproduced for motor skills that spontaneous α-band FC of the motor ROI with the rest of the brain correlated with performance (the average number of completed sequences per minute) of the participants (Fig. 8 C, r=0.48, p=0.033). This was again not the case for FC in other frequency bands.

**Figure 8.**
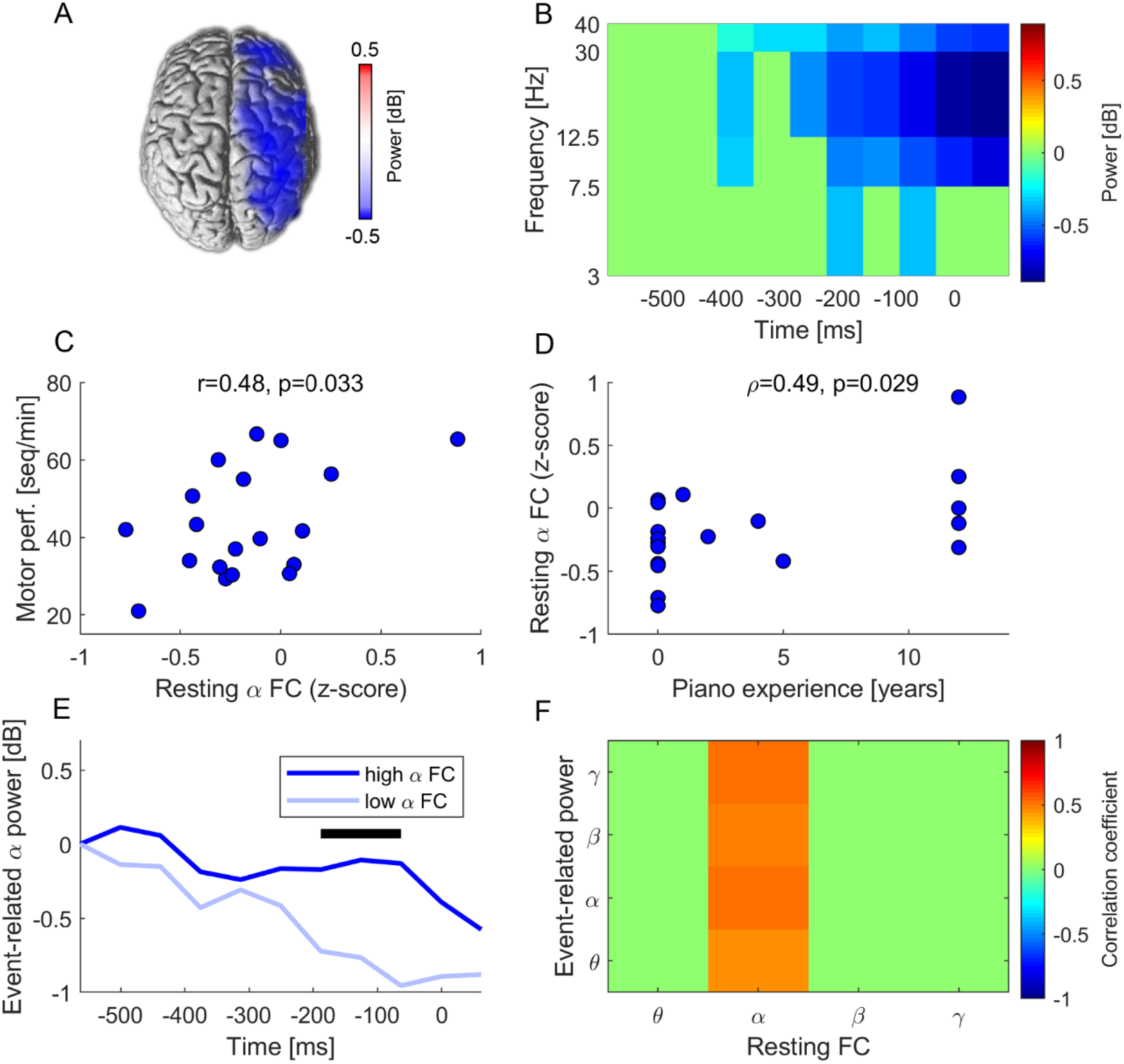
Generalization to motor planning. Mean event-related α-band power decrease before execution of a motor sequence across all 20 patients (p<0.05, 5% FDR corrected) (**A**). A time-frequency decomposition at the contralateral motor ROI shows, on average, prominent power decrease in all examined bands (p<0.05, 5% FDR corrected). Time 0 indicates the onset cue (**B**). Correlation between spontaneous α-band FC and number of correct motor sequences per minute (**C**). Greater α-band FC was in turn associated with more years of experience with piano playing (**D**). Participants with low spontaneous α-band FC needed significantly greater event-related α power decrease than subjects with high FC (**E**). Black line indicates time points with significant difference. Thus, greater spontaneous α-band FC correlated with proportionally smaller task-induced power decrease at all examined frequency bands (**F**, p<0.05, corrected).

Crucially, α-band FC, but not FC in the other frequency bands, correlated with the number of years of piano playing (Fig. 8 D), hence suggesting that previous experience with the task was linked to the appearance of abundant α-band coupling. Piano experience was also positively correlated with performance (ρ=0.77, p<0.0001).

When dividing the subjects into two equal subgroups of high and low spontaneous α-band FC (median cutoff), we again observed that subjects with low spontaneous FC needed greater power decrease in θ (F_1,189_=9.4, p=0.0025), α (F_1,189_=18.1, p<0.0001), and β bands (F_1,189_=12.2, p=0.0006) than participants with high FC (Fig. 8 E). Furthermore, the amount of spontaneous α-band FC correlated with the area under curve of ERDs in all examined frequency bands (Fig. 8 F), similarly as in experiment 1.

## Discussion

We reproduced, on average, the classical local power decrease (i.e., ERD) which is widely used as an index of cortical activation (Pfurtscheller and Lopes da Silva, 1999). At the single trial level, ERDs had a positive, independent impact on perception in a within-subject analysis. However, this result was driven by only part of the sample, as some subjects show little to no ERDs following target presentation, whether that target was seen or missed. Furthermore, if post-target ERD is indeed involved in perception or action planning in some subjects, our results show that it is not the most efficient processing pathway. Indeed, regarding inter-individual differences in visual perception performance, we uncovered that a greater in-task power decrease in seen than missed trial was actually associated with *worse* performance. Moreover, the subjects who were able to detect the highest percentage of targets of a given contrast, or to complete the greatest number of motor sequences, were those in which the event-related power decrease was the smallest, tending towards an absence of difference.

We also reproduce previous findings that states of higher network coupling enable efficient neural processing and high performance (Hipp *et al*., 2011*a*; Weisz *et al*., 2014; Sadaghiani *et al*., 2015; Strauß *et al*., 2015). However, most previous research has still used classical paradigms with externally induced task repetitions and has thus essentially left untouched the classical primacy of momentary and local task-induced activations.

Our article compares the impact of *spontaneous* functional connectivity states across much longer time scales to the impact of classical task-induced brain activations on behavior. The critical novelty is thus that spontaneous functional connectivity states expressed over several seconds, in particular α band phase coupling, are the primary correlate of task success. In contrast, activations induced by classical task contexts are minimal or absent in high performers, which demonstrates that they are merely a compensation mechanism needed only in participants with unfavorable connectivity states. Furthermore, the level of functional coupling predicted the ERD magnitudes in a wide range of frequency bands. Our findings thus show that ERDs can at best be seen as index of effortful attention (Wyart and Tallon-Baudry, 2008) which enables low-performing subjects to accomplish a task despite unfavorable neural states. In contrast, intrinsic properties of the network, in particular their coupling states, enable more efficient processing without the need for local activations. This may reflect the underlying synaptic architecture with pre-established pathways for distributed processing of the task (Weisz *et al*., 2014). This also means that network states need to be taken into account in any attempt to explain inter-individual differences in behavior and task-related neural events (Iemi *et al*., 2019).

Evidence for a causal role of spontaneous coupling on performance comes from neurofeedback studies demonstrating that enhancing neural coupling improves behavioral performance in healthy (Kajal *et al*., 2017; Koush *et al*., 2017) and in patients with brain lesions (Mottaz *et al*., 2018). This also means that spontaneous neural coupling may become a primary target in programs that aim to improve performance. Indeed, experiment 2 shows that long-term experience with a task associates with greater spontaneous neural coupling of the involved brain areas. Future work will need to examine which training strategies are most efficient for enhancing spontaneous neural coupling. In particular, it may be possible to use complementary and more efficient methods to classical repetitive task training.

ERDs found after unilateral stimulation in experiment 1 were bilateral. Conversely, activity induced by unilateral targets is classically shown to be contralateral to their presentation. However, several aspects of our paradigm may explain this. First, we provided no cueing on the location of the next target. The best strategy was thus to attend to both locations for all trials. A consistent shift would be expected to cause at least a small modulation of threshold contrast (Wyart and Tallon-Baudry, 2008). As overall performances did not differ from 50% accuracy, we can infer that they maintained a bilaterally distributed focus throughout the titration and recording runs. Second, the fact that targets could be presented bilaterally diminishes certainty even after one target has been perceived. In the case of spatial cueing, the level of validity of the cue (i.e. yielding a given spatial certainty) influences the degree of α-power lateralization (Rohenkohl and Nobre, 2011). We suggest that a similar phenomenon occurs in our task: following unilateral targets, there is a shift of attention toward the side. But because there is uncertainty about the simultaneous occurrence of a second target to the other side, the effect on α-power is less lateralized than in paradigm that include only one-sided stimulations.

High-γ power increase is, as well as α ERD, predictive of perceptual decisions (Wyart and Tallon-Baudry, 2009). Furthermore, high-γ activations are usually more circumscribed and unilateral and more task-specific (Crone *et al*., 2006). Figures 6 and S4 demonstrate that our conclusions also apply to high-γ activations.

New models of brain functioning are required to integrate our observations. The importance of network interactions for conscious processing, in particular for visual consciousness is already well grounded in connectivist models such as the global neuronal workspace theory of consciousness (Dehaene and Changeux, 2011). However, our findings additionally demonstrate the importance of spontaneous states outside of the task context.

How might spontaneous network states influence behavior and determine the need for task-induced activations? The neural efficiency theory (Haier *et al*., 1988) suggests that higher intelligence is underpinned by a higher efficiency in its functioning rather than a higher excitability or α-band ERD as an index of cortical activation (Grabner *et al*., 2004). This framework has been extended beyond the notion of intelligence to expertise: skilled populations (e.g., expert chess players or athletes) have a reduction in task-induced brain activation as compared to naïve subjects (Milton *et al*., 2007; Percio *et al*., 2010), in accordance with our findings from experiment 2. Recent research identified neuronal coupling as underlying electrophysiological correlate of neural efficiency and expertise. Network interactions were found to be stronger in expert brains at rest (Gong *et al*., 2019) and during tasks related to their field of expertise (Bhattacharya and Petsche, 2005). In this view, α-band coupling thus reflects general intelligence and expertise.

The predictive processing theory goes a step further in suggesting that expertise arises from a refined model of the self and world which allows to make more accurate predictions about upcoming events (Friston and Stephan, 2007; Euler, 2018). In this framework, network interaction states could be seen as reflecting expectancies (Mayer *et al*., 2016) which help reduce the indeterminacy of visual stimuli or motor sequences. This could in turn result in reduced ERDs reflecting less prediction error, i.e., less difference between expected and observed events. Although we did not modulate indeterminacy in our paradigms, it has been argued that past experience, as in our musicians of experiment 2, will make low contrast visual stimuli or motor sequences less novel and indeterminate (Euler, 2018).

It is interesting to note that the brain has a preference for α frequencies for long-range neural interactions. We underline that it is spontaneous α-band coupling that predicted performance, whereas local α power did not. Therefore, the frequency-specificity cannot be to a higher signal-to-noise ratio of α activity. Rather α seems to have a primary role in neural communication during rest which we show here to enable high task performance. Most previous propositions on the role of α activity have been derived from classical task-related experiments, which is precisely the situation in which α is replaced by faster rhythms. The idea of an inhibitory role may thus be an artifact resulting from the experimental context (Palva and Palva, 2011).

Besides the α-frequency that was of primary interest here, we also observe that phase coupling in γ frequencies influences ERD in visual areas (Fig. 5 H). This matches well with animal studies: intra-cortical recordings in monkey visual cortex have shown that α exerts a top-down modulation on visual processing, through its influence on γ activity. Within V1, these two rhythms display phase-amplitude coupling as well as anti-correlated power (Spaak *et al*., 2012). Γ oscillations travel downstream from primary (V1) to secondary (V4) areas, whereas α oscillations flow upstream (Van Kerkoerle *et al*., 2014). In relation with our results, we can thus hypothesize that a high sensitivity to feedback from higher level to primary areas, as evidenced by a high level of FC in the α band, is an advantage for stimulus detection.

## Methods

### Participants

Twenty healthy subjects were analyzed for experiment 1 (12 women; 27.9±4.9 years old) and twenty for experiment 2 (13 women; 28.7±5.6 years old). Two additional subjects of experiment 1 were excluded from analysis due to poor quality of the EEG recording. All had normal or corrected-to-normal vision and no history of neurological or psychiatric disorders and were paid for their participation.

All procedures were approved by the ethical committee of the canton of Geneva and performed according to the declaration of Helsinki. All participants gave written informed consent after receiving explanation on the nature the experiments.

### Experimental paradigm and stimulus presentation

Subjects were seated 54 cm away from a 23.5’’ Eizo FG2421 LCD monitor with a refresh rate of 120 Hz.

<u>Experiment 1</u>: Stimulus presentation was implemented in MATLAB (The MathWorks, Natick, USA) using the Psychophysics Toolbox extensions (Brainard, 1997). The experimental paradigm is illustrated in Figure 2. Participants were cued to close their eyes by an instruction screen. The resting period lasted between 15 and 20 s. An auditory cue signaled the end of the resting period and the imminent start of a series of trials. Participants opened their eyes and kept fixation on a central cross (0.5° visual angle) to avoid eye movement throughout the time of each trial. Two light grey squares (3×3°) were displayed on a darker grey background and served as position markers for the location of target presentation. They were centered at 8° vertical and 24° horizontal eccentricity from the fixation cross. After a delay of 1500 ms, targets (circular patches of 0.5° visual angle with a Gaussian envelope) were displayed for 42 ms (5 time frames) at perithreshold contrast in the center of the position markers, either on the left, on the right, or bilaterally. Each block comprised two occurrences of each experimental condition (left, right and bilateral) and one catch trial in which no target was displayed. The order of presentation was randomized. After another 1500 ms, a response screen was displayed and participants had to answer, for each side, whether they had seen a target using 4 different response buttons (two for each side). There was no time limit for response which could be made simultaneously for both sides or in any sequential order. They were instructed to avoid guessing and only give a positive answer when they were confident that a target had been displayed. The next trial began as soon as an answer had been collected for each side.

<u>Experiment 2</u>: The sequential finger tapping task (FTT) was designed using E-Prime 2.0 software (Psychology Software Tools, Pittsburgh, PA). Participants were instructed to repeat a given five-item sequence with their left hand (little finger to index) on four horizontally arranged buttons numbered left to right on a Chronos box (Psychology Software Tools, Pittsburgh, PA; https://pstnet.com/products/chronos/). The same sequence was trained throughout the whole experiment (1-4-2-3-1). It was continuously presented to participants while they had to perform it. During test blocks obtained for assessment of performance, they had to repeat the sequence as fast and accurately as possible during 30 seconds. No feedback was given. A further block with EEG recording was obtained to reconstruct neural activations during the task. Each trial of the EEG block was composed of two consecutive sequences with a delay of 1.5 seconds between trials (end to beginning), cued with the apparition of the sequence on the screen, for a total of 150 trials. Feedback was given with a star appearing under each item correctly pressed. In case of error, participants had to repeat the item until correct.

### Procedure

<u>Experiment 1</u>: Subjects were tested over two consecutive afternoons. On day one, titration runs were obtained to determine individual perithreshold contrasts for each subject and condition. The titration session comprised two runs. The first run consisted in a 1-up 1-down interlaced triple staircase procedure with a decreasing step size and provided a rough estimate of the threshold contrast for each condition. The second run aimed to refine the thresholds and was based on the method of constant stimuli: 6 levels of contrast were chosen for each condition based on the point of convergence of the staircase and presented 16 times each in a random order, enabling us to estimate the psychometric function for each subject and condition (Treutwein, 1995) and thus to choose contrast values with a 50% probability of detection. For bilateral trials, this meant values resulting in a 50% probability of both targets being simultaneously detected. Awareness of only one (either left or right) or none of the targets in a bilateral trial was considered an incomplete/incorrect detection. In both runs the three experimental conditions (left/right/bilateral) and catch trials (no target) were presented in a random order.

The EEG data were recorded during day two. A first (shorter) run was performed to check that the subjects detected approximately 50% of the targets. It also permitted to familiarize them with the resting part of the task. For 15 subjects, the level of contrast had to be adjusted (max. 0.3% increase/decrease) for a least one condition. Participants then completed 120 blocks of the task, divided in 6 sessions of 20 blocks.

<u>Experiment 2</u>: The experiment consisted of one session. Participants completed test blocks before and after the EEG block. Participants could take a short break every 50 trials. Eyes-closed resting-state EEG was recorded during a 10 minutes before the first test block, task-induced EEG during the EEG block.

### Behavioral analyses

<u>Experiment 1</u>: To assess compliance with instructions – participants were asked to adopt conservative criteria – we examined false alarm rates and further computed Grier’s B’’, a nonparametric measure of response bias that ranges from -1 (maximum bias towards *yes* answers) to +1 (maximum bias towards *no* answers) (Grier, 1971; Stanislaw and Todorov, 1999). One-sample t-tests were used to check whether the chosen contrast values elicited a performance of approximately 50% accuracy in our three conditions. We compared the threshold contrast values (after adjustment on recording day) between conditions using a Friedman test, given the non-normal distribution of this variable in all conditions (Shapiro-Francia tests, all p<.03). In order to obtain an estimate of visual performance comparable across subjects, we computed the mean detection rate across conditions for contrasts that were presented to all subjects during the second titration run. Better visual performance results in a higher visual perception percentage. To assess visuo-spatial bias, we calculated the difference between the number of targets detected on the left versus on the right for bilateral trials administered on day two. A greater bias score is found in subjects who detected the left target more often than the right.

<u>Experiment 2</u>: Performance was assessed as the mean number of correctly completed sequences per minute in all test blocks.

### EEG acquisition

EEG data were sampled at 1024 Hz using a 128-channel BioSemi ActiveTwo EEG-system (BioSemi B.V., Amsterdam, Netherlands). Data were re-referenced against the Cz electrode. Artifacts such as eye movements, blinks, power line, and electrode artifacts were removed by visual inspection in experiment 1 and for resting data of experiment 2, and using independent components analysis (FastICA algorithm (Hyvarinen, 1999)) for task-induced epochs in experiment 2.

### Source imaging

All analyses were performed in MATLAB (The MathWorks, Natick, USA), using the toolbox NUTMEG (Dalal *et al*., 2011) and its Functional Connectivity Mapping (FCM) toolbox (Guggisberg *et al*., 2011).

The α frequency band (8-12 Hz) was the primary band of interest. Other frequency bands (θ, β, γ, high-γ; respectively 3-7, 13-30, 31-45 and 55-100 Hz) were included to assess band-specificity. The high-γ band was only examined in experiment 1, as the ICA pre-processing of experiment 2 needed to be limited to 1 to 40 Hz.

Lead-potential was computed using a boundary element head model, with the Helskini BEM library (Stenroos *et al*., 2007) and the NUTEEG plugin of NUTMEG (Guggisberg *et al*., 2011). The head model was based on the individual T1 MRI of each participant and solution points were defined in the gray matter with 10 mm grid spacing. EEG data was bandpass-filtered in the respective frequency bands, Hanning windowed, Fourier transformed and projected to grey matter voxels using an adaptive filter (scalar minimum variance beamformer) (Sekihara *et al*., 2004). Regions of interest (ROIs) were defined anatomically. We chose the dorsal V3 region (Eickhoff *et al*., 2005; Kujovic *et al*., 2013) for experiment 1 since this region specifically responds to stimulation of the lower quadrant and V3 neurons have been shown to have very low contrast thresholds (Gegenfurtner *et al*., 1997; Larsson and Heeger, 2006). Moreover, EEG coverage is better for dorsal parts of the visual cortex. For experiment 2, a ROI combining M1 and the dorsal premotor area contralateral to the moved arm was used (Mayka *et al*., 2006). Analysis of FC was conducted as described previously (Guggisberg *et al*., 2011, 2015). We used the imaginary component of coherence (IC) as index of FC and calculated the weighted node degree (WND) for each voxel as the sum of its coherence with all other cortical voxel (Newman, 2004). The artifact-free resting-state data was segmented into non-overlapping 1s epochs.

<u>Experiment 1</u>: To compute overall levels of spontaneous FC, 1500 artifact-free resting-state epochs were randomly selected for each participant. To examine variations of FC in and their influence on subsequent performance, we calculated FC values for each block, based on the last twelve seconds of each resting period.

Task-induced power modulations were reconstructed from 500 ms before to 750 ms after target onset using the same pass-bands and inverse solutions as for FC. Single trial power at the visual ROIs was computed as the average root-mean square of source time series across ROI voxels for a total of 176 ± 29 epochs for each subject. Average values across trials were computed after baseline subtraction for each participant. In addition, mean power differences between seen and missed trials were obtained from non-baseline corrected values to avoid confounds arising from baseline differences. They were computed as area under curve of the whole peri-stimulus period. For voxel-wise imaging, a 200ms long sliding window with a 50ms time step was used.

<u>Experiment 2</u>: Spontaneous FC was obtained from 300 artifact-free epochs of 1 s recorded during rest.

Task-induced power modulations were reconstructed from 562.5 ms before to 62.5 ms after the sequence cue, with 125ms long sliding windows and 62.5ms steps using 1-40Hz band-pass data. Average power across 100 artifact-free epochs was obtained for each participant. In addition the area under curve of power change was obtained for time windows with significant differences between high and low FC participants.

## Statistical analyses

Power values were log-transformed to obtain normal distributions.

To determine whether variations of our electrophysiological measures influence perception of subjects in experiment 1, one-sample t-tests were applied on the difference in FC and in event-related power between seen and missed trials. Single trial data was subjected to mixed-model ANOVAs to assess multivariate effects of FC and in-task power on perception.

Between-subject Pearson correlations were used to test the relations between subject’s mean values of performance, resting-state FC and the area under curve of in-task power decreases, for data from both experiments. No outliers were present that could have biased the correlation estimates. To further explore those relations, we split the subject sample in halves using the median resting-state α- band FC level at the ROI as group cutoff criteria (low versus high). We then compared event-related power in high vs. low FC subjects using repeated-measures ANOVAs. In experiment 1, we also examined the effect of perception and time on in-task power with repeated-measures ANOVAs.

Correction for multiple comparisons was achieved through a 5% false discovery rate.

## Resource Availability

NUTMEG, the Matlab toolbox used for these analyses is available at http://nutmeg.berkeley.edu.

## Author contributions

LA performed and analyzed experiment 1 and wrote the paper. AM performed and analyzed experiment 2 and assisted in experiment 1. AK supervised the project. AGG designed and supervised the experiments and wrote the paper.

