## Supplemental figures 1 to 4 for "Spontaneous network coupling enables efficient task performance without local task-induced activations"

### Supplemental information

#### Supplemental Figures

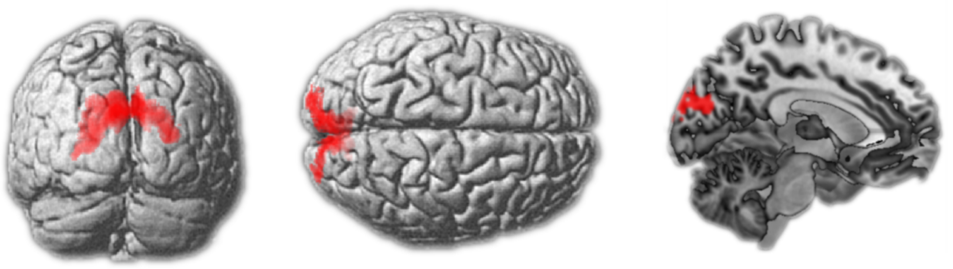

**Figure S1. Occipital region of interest.** Dorsal V3 region anatomically defined from the Jülich probabilistic cytoarchitectonic atlas (Eickhoff *et al.*, 2005; Kujovic *et al.*, 2013).

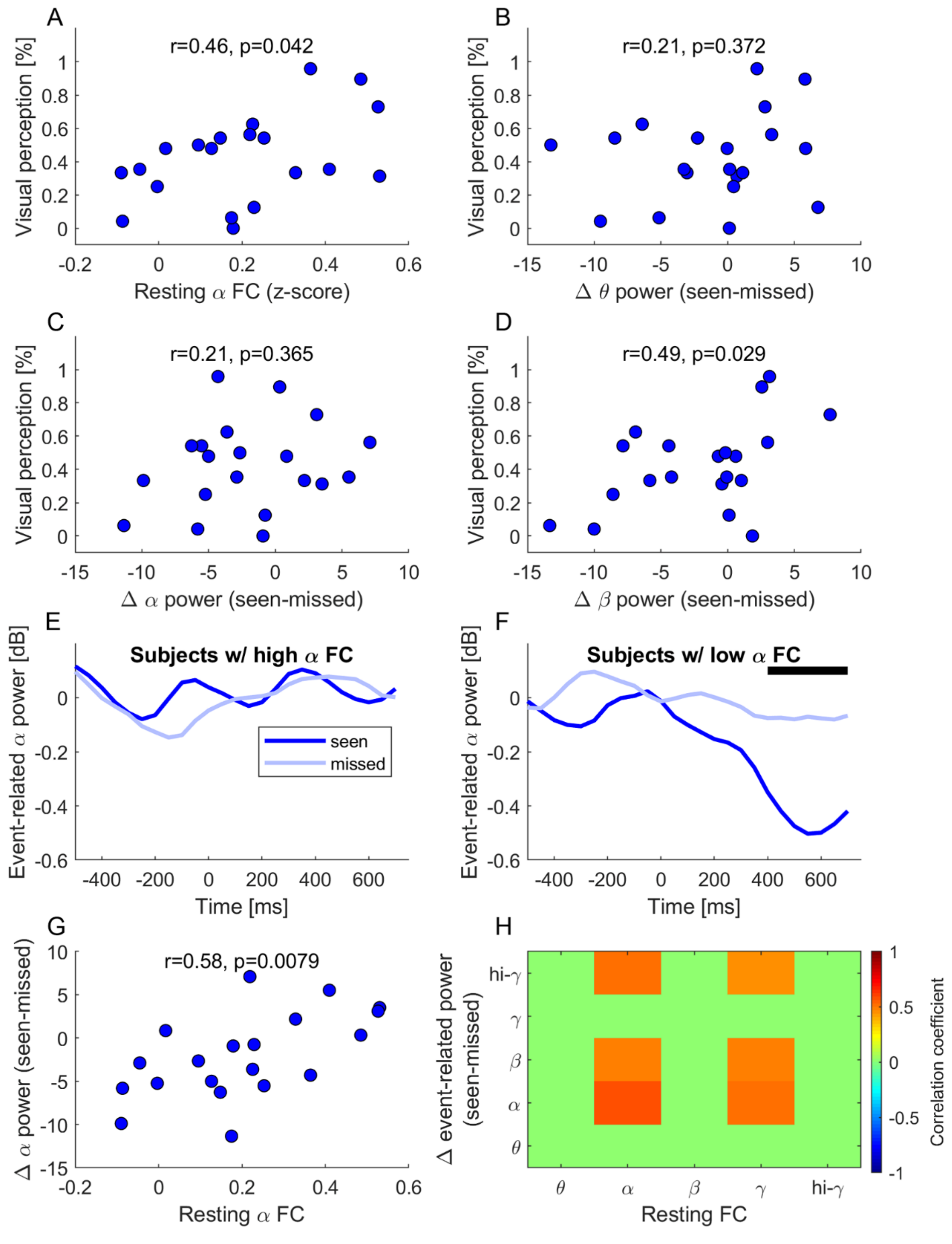


**Figure S2. Between-subject co-variation of neural processing and target perception; left targets.** Association between resting-state α WND and mean detection rate at optimal contrast (**A**). Association between in-task power difference (seen-missed) in θ, α, and β bands and mean detection rate at optimal contrast (**B-D**). α-band power variation over the peri-stimulus interval for seen and missed targets in the ten subjects with the lowest (**E**) or highest (**F**) resting-state α WND. The black line indicates time windows in which the difference between seen and missed trials is significant ($p < 0.05$). Association between resting-state α WND and in-task α power difference (**G**) and same association for all bands (**H**).

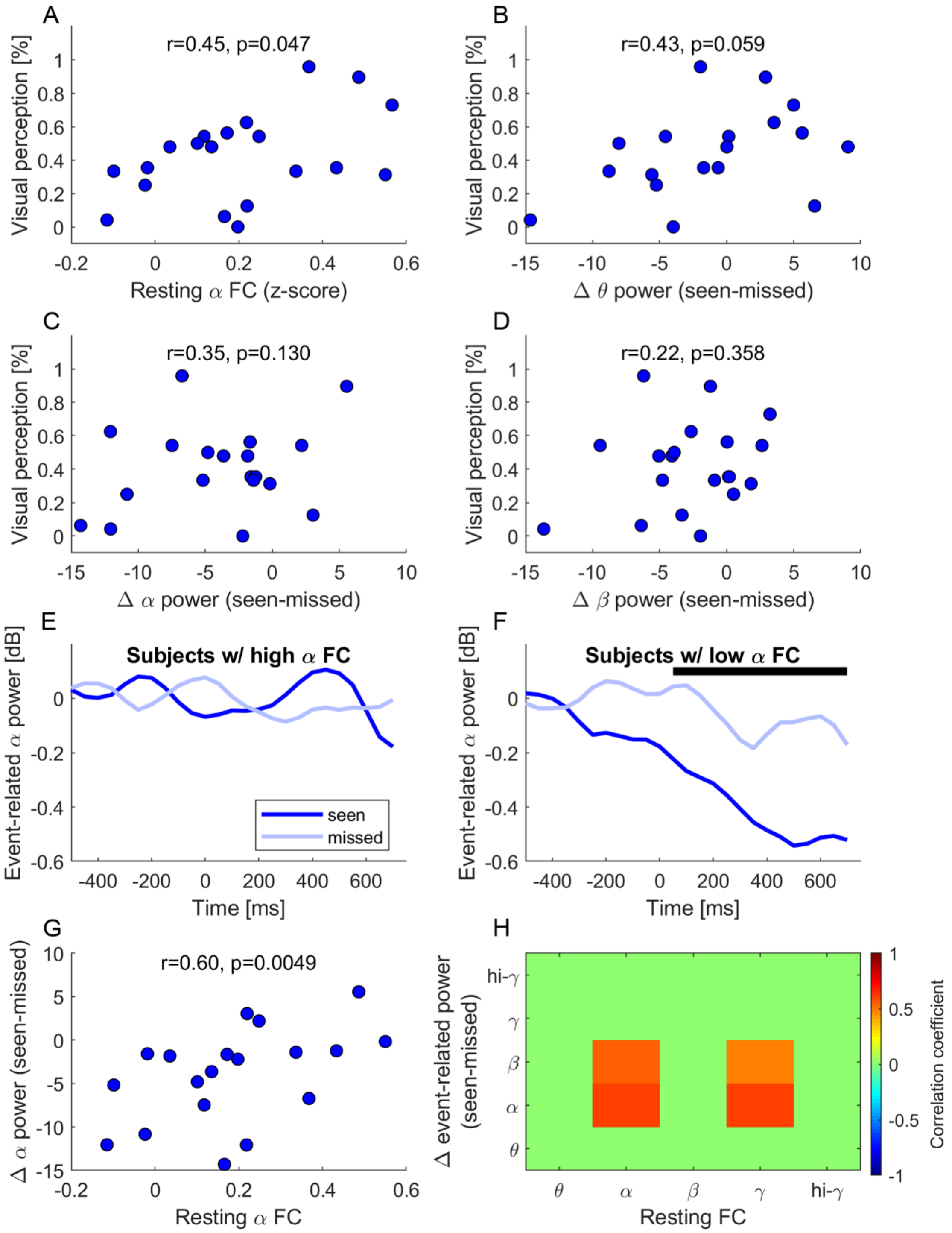


**Figure S3. Between-subject co-variation of neural processing and target perception; right targets.** Association between resting-state α WND and mean detection rate at optimal contrast (**A**). Association between in-task power difference (seen-missed) in θ, α, and β bands and mean detection rate at optimal contrast (**B-D**). α-band power variation over the peri-stimulus interval for seen and missed targets in the ten subjects with the lowest (**E**) or highest (**F**) resting-state α WND. The black line indicates time windows in which the difference between seen and missed trials is significant ($p < 0.05$). Association between resting-state α WND and in-task α power difference (**G**) and same association for all bands (**H**).

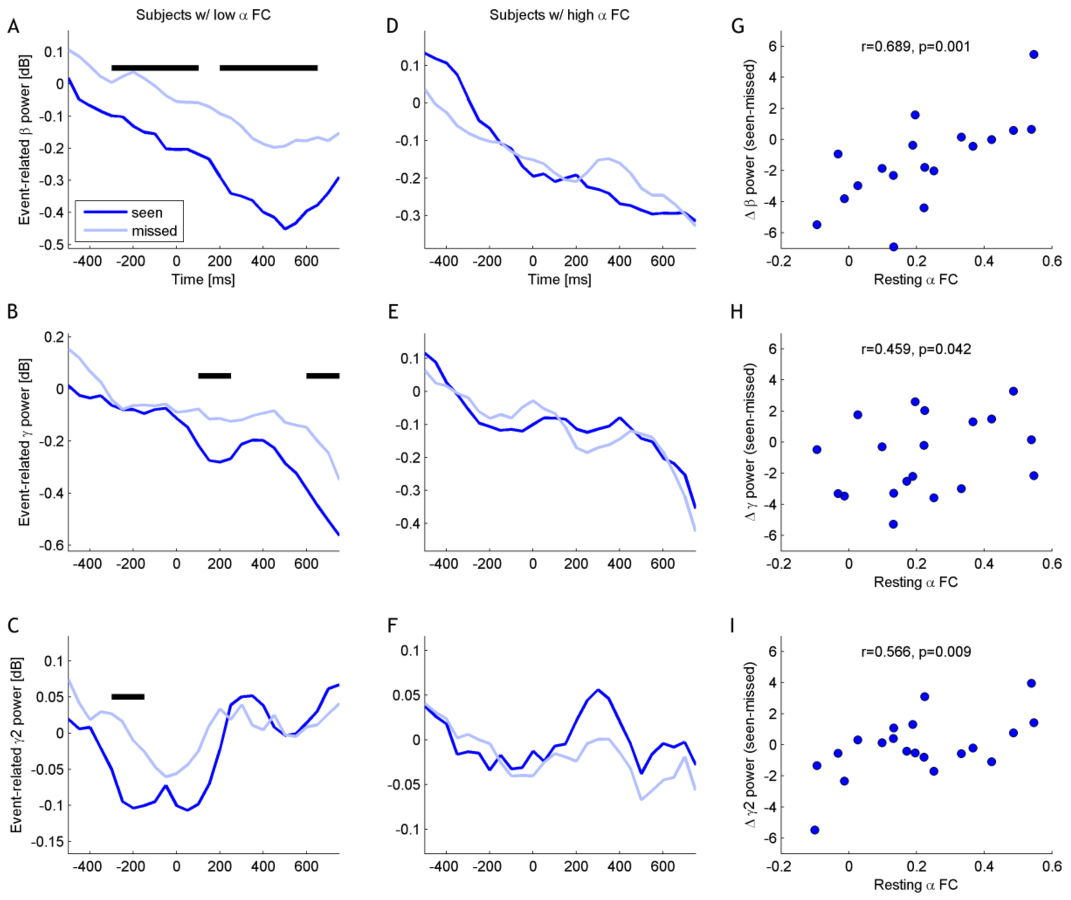


**Figure S4.** Power variation over the peri-stimulus interval for seen and missed targets in $\beta$, $\gamma$ and high-$\gamma$ in the ten subjects with the lowest (**A**, **B**, **C**) or highest (**D**, **E**; **F**) resting-state $\alpha$ WND. The black line indicates time windows in which the difference between seen and missed trials is significant ($p < 0.05$). Association between resting-state $\alpha$ WND and in-task power difference in $\beta$ (**G**), $\gamma$ (**H**) and high-$\gamma$ ($\gamma 2$) frequency bands (**I**; Pearson correlations).
